# Microbiomes of hadal fishes contain similar taxa, obligate symbionts, and known piezophiles across trench habitats

**DOI:** 10.1101/2022.01.15.476391

**Authors:** Jessica M. Blanton, Logan M. Peoples, Mackenzie E. Gerringer, Caroline M. Iacuniello, Natalya D. Gallo, Thomas D. Linley, Alan J. Jamieson, Jeffrey C. Drazen, Douglas H. Bartlett, Eric E. Allen

## Abstract

Hadal snailfishes are the deepest-living fishes in the ocean, inhabiting trenches from depths of ∼6,000 to 8,000 m. While the microbial communities in trench environments have begun to be characterized, the microbes associated with hadal megafauna remain relatively unknown. Here, we describe the gut microbiomes of two hadal snailfishes, *Pseudoliparis swirei* (Mariana Trench) and *Notoliparis kermadecensis* (Kermadec Trench) using 16S rRNA gene amplicon sequencing. We contextualize these microbiomes with comparisons to the abyssal macrourid *Coryphaenoides yaquinae* and the continental shelf-dwelling snailfish *Careproctus melanurus*. The microbial communities of the hadal snailfishes were distinct from their shallower counterparts and were dominated by the same sequences related to the Mycoplasmataceae and Desulfovibrionaceae. These shared taxa indicate that symbiont lineages may have remained similar to the ancestral symbiont since their geographic separation or that they are dispersed between geographically distant trenches and subsequently colonize specific hosts. The abyssal and hadal fishes contained sequences related to known, cultured piezophiles, microbes that grow optimally under high hydrostatic pressure, including *Psychromonas, Moritella*, and *Shewanella*. These taxa are adept at colonizing nutrient-rich environments present in the deep ocean, such as on particles and in the guts of hosts, and we hypothesize they could make a dietary contribution to deep-sea fishes by degrading chitin and producing fatty acids. We characterize the gut microbiota within some of the deepest fishes to provide new insight into the diversity and distribution of host-associated microbial taxa and the potential of these animals, and the microbes they harbor, for understanding adaptation to deep-sea habitats.

**Importance:** Hadal trenches, characterized by high hydrostatic pressures and low temperatures, are one of the most extreme environments on our planet. By examining the microbiome of abyssal and hadal fishes, we provide insight into both the physiology of the deepest-living vertebrates and the microbes which colonize them. Our findings show that there are similar microbial populations in fishes geographically separated by thousands of miles, reflecting strong selection for specific microbial lineages. Only a handful of psychropiezophilic taxa, which do not reflect the diversity of microbial life at great depth, have been successfully isolated in the laboratory. Our examination of deep-sea fish microbiomes shows that typical high-pressure culturing methodologies, which have largely remained unchanged since the pioneering work of Claude ZoBell in the 1950s, may simulate the chemical environment found in animal guts and helps explain why the same deep-sea genera are consistently isolated.

## Introduction

The gut microbiome plays an essential role in the physiology of fishes. Microbiota within fishes can help digest food by producing degradative enzymes, provide the host with vitamins and fatty acids, and competitively exclude pathogens (1, 2, 3, 4). While the importance of gut microbiomes is recognized, few studies have explored the structure and function of microbiomes in deep-sea fishes. Cultivation of microorganisms from deep-sea animals has revealed the presence of piezophiles (5, 6, 7, 8), microbes capable of optimal growth under *in situ*, deep-sea high hydrostatic pressure conditions. This includes members of the genera *Colwellia, Psychromonas, Shewanella, Moritella*, and *Photobacterium*, some of the only lineages which have been experimentally demonstrated in the laboratory to be piezophilic (9, 10). These microbes represent a small fraction of the broader water and sediment communities in the deep ocean, which are instead composed primarily of members of the Thaumarchaeota, Marinimicrobia (SAR406), and other members of the Proteobacteria (11, 12, 13, 14). However, a description of the complete breadth of microbial diversity within the guts of deep-sea fishes is lacking.

Distinct fish communities have evolved to life in the deep sea, with pronounced compositional changes within different depth zones (15). The abyssal ocean (depths 4,000–6,000 m) is home to several major fish families with cosmopolitan distributions, including the rattails (Macrouridae), cusk eels (Ophidiidae), eelpouts (Zoarcidae), cutthroat eels (Synaphobranchidae), and tripodfishes (Ipnopidae). Rattails are attracted to bait and therefore have been the focus of much of the deep-sea demersal fish literature. Members of the rattail genus *Coryphaenoides*, which includes *Coryphaenoides yaquinae* Iwamoto and Stein 1974 (16) and *Coryphaenoides armatus* Hector 1875 (17), are among the most widespread fishes in abyssal ecosystems (18). *Coryphaenoides* species are known scavengers (19, 20, 21, 22) and their predominant food source is deep-sea carrion, although stomach contents and stable isotope analyses show that rattails also feed on fishes, squid, and crustaceans (18). Culture-based analyses of the microbiota associated with *Coryphaenoides* have found piezophilic members related to the lineages *Moritella* and *Shewanella* (5, 6). However, whether these lineages are representative of the entire microbiota within the gut of *Coryphaenoides*, one of the most widespread fishes in the ocean, is unknown.

In hadal trenches, sites deeper than 6,000 m which are typically formed at subduction zones, the fish community differs from that of the surrounding abyssal plain. Snailfishes (family Liparidae) are the dominant fishes below 6,000 m, with at least twelve species found in nine trenches worldwide (23). The Liparidae include the planet’s deepest known vertebrates, such as *Pseudoliparis swirei* Gerringer & Linley 2017 (24; depth range 6198–8078 m) and *Notoliparis kermadecensis* Nielsen 1964 (24, 25, 26; depth range 5879–7669 m). Many hadal snailfish species have been found in only one trench and are likely endemic, confined to one specific hadal environment (23, 25, 26, 27). These fishes have evolved adaptations to high pressure, including intrinsic enzyme adaptations (28, 29) and the accumulation of protein-stabilizing osmolytes such as trimethylamine n-oxide (TMAO; 30, 31). No fishes have been found deeper than ∼8,200 m, a putative physiological depth limit for vertebrates arising from the osmotic constraints of this TMAO pressure-adaptation strategy. Stomach contents, stable isotope analyses, and observed feeding behavior indicate that snailfishes are one of the top predators at hadal depths, consuming highly-abundant amphipods in trench habitats (26, 32, 33). Recently, a member of the Mycoplasmataceae was identified in *Pseudoliparis swirei* that may provide the host with riboflavin (34). However, it is unclear how differences in fish species, diet, and environmental conditions may influence the composition of gut microbiomes of abyssal and hadal fishes, or how these microbial associates may impact the physiology of the host.

Here, we describe the gut microbiota of four representative, ecologically-important deep-sea fishes using 16S rRNA gene amplicon sequencing. This includes two of the deepest-living hadal snailfishes, *Pseudoliparis swirei* from the Mariana Trench and *Notoliparis kermadecensis* from the Kermadec Trench. These fishes and the trenches they inhabit are geographically separated, residing approximately 6,000 km apart within the Pacific Ocean. The Mariana Trench is located in the Northern Hemisphere and extends to a depth in excess of 10,900 m (35). The Kermadec Trench is in the Southern Hemisphere off the coast of New Zealand and reaches a depth exceeding 10,000 m (36). We compared the microbiota of the snailfishes with two shallower-dwelling fishes, the abyssal macrourid *Coryphaenoides yaquinae* which inhabits depths of ∼3,000 to 7,000 m (26), and *Careproctus melanurus* Gilbert 1892 (37), a demersal snailfish typically found at depths 200 – 1,600 m (38). Our findings inform new understanding of host-symbiont interactions in the abyssal and hadal ocean, the ecology of piezophilic microbes, and the biology of the planet’s deepest-living vertebrates.

## Results

The gut microbial communities within snailfish from the Mariana Trench (*Pseudoliparis swirei*; n=18, collection depths 6,898 – 7,966 m) and Kermadec Trench (*Notoliparis kermadecensis;* n=7, collection depths 6,456 – 7,515 m) were compared against those in a continental shelf-dwelling snailfish (*Careproctus melanurus*; n=11, collection depths 381 – 834 m) and an abyssal rattail (*Coryphaenoides yaquinae*; n=4, collection depths 4,441 – 6,081 m; **Figure 1**; **Table S1**). We identified a total of 2,034 Amplified Sequence Variants (ASVs) across these four species with final amplicon libraries ranging from 2,545 – 106,059 reads per sample (average ∼46,500 reads per sample).

**Figure 1.**
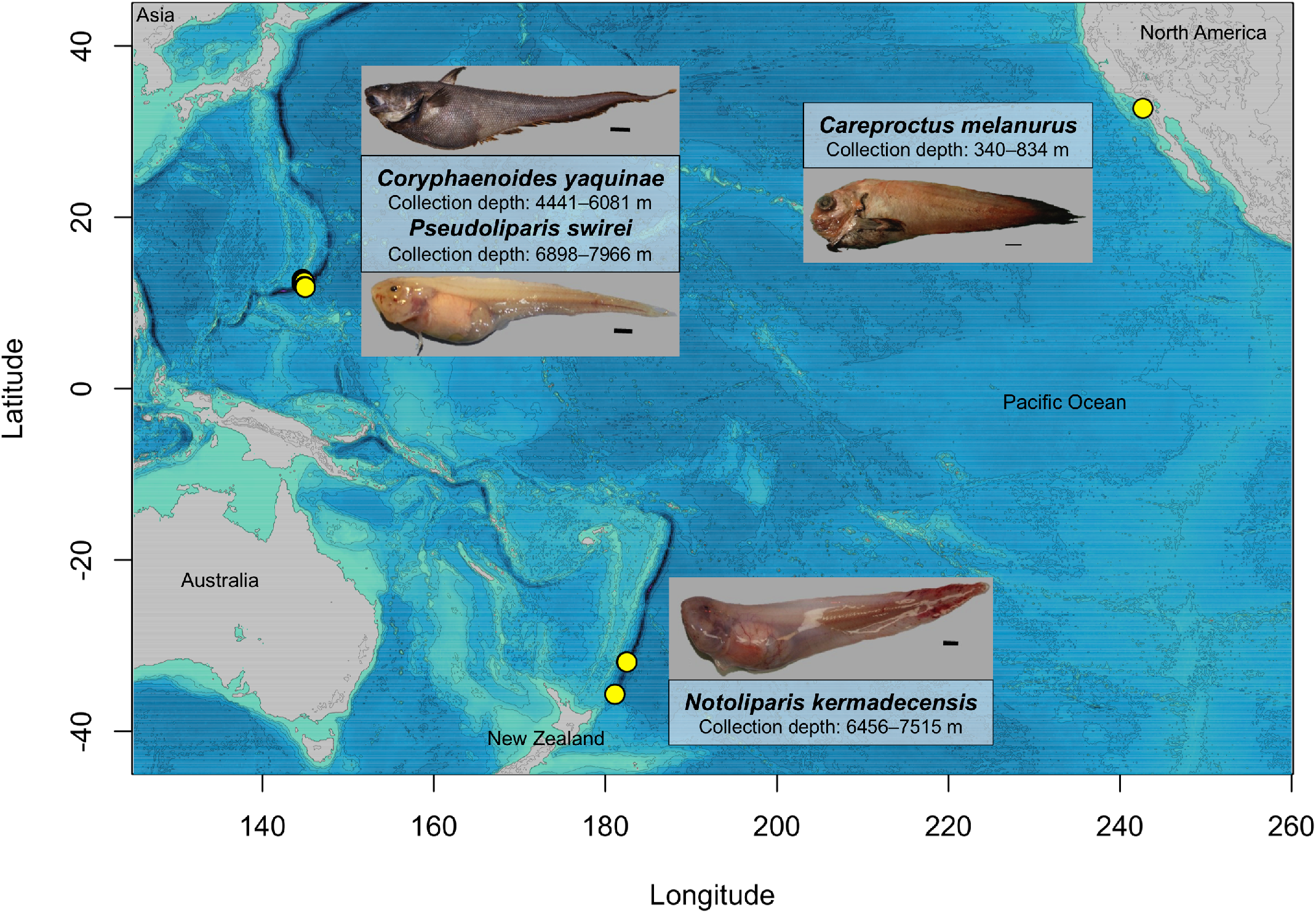
Map of the Pacific Ocean showing the locations and depths of collection of the four fish species described in this study. a) *Pseudoliparis swirei* (n=18), Mariana Trench, 7626 m, #200133, scale bar 1 cm. b) *Notoliparis kermadecensis* (n=7), Kermadec Trench, 7515 m, #100171, scale bar 1 cm. c) *Coryphaenoides yaquinae* (n=4), abyssal plain, 5255 m, #200152, scale bar 5 cm. d) *Careproctus melanurus* (n=11), continental slope, representative image, scale bar 1 cm.

### Fish microbiome comparative analyses

Gut microbial communities were distinct between the hadal snailfishes, the rattail *C. yaquinae*, and the slope-dwelling *C. melanurus*. The hadal fishes had lower alpha diversity than the shallower fishes, with the gut microbiome of *C. yaquinae* appearing more even (**Figure 2, Figure S1)**. NMDS ordination analysis of Bray-Curtis dissimilarity demonstrated that microbial gut communities of each fish species were distinct from one another, where species type accounted for 37% of the variability (**Figure 2C**; PERMANOVA, R^2^ = 0.37, F = 7.22, DF = 3, p < 0.001). Pairwise comparisons showed that while the microbiome of *Pseudoliparis swirei* differed from that in *Notoliparis kermadecensis* (R^2^ = 0.09, p < .013, F = 2.52), these differences were small in comparison to the other fishes. For further context, the gut microbiomes of the four species of interest were compared to those from a diverse collection of fishes. This dataset included 16 marine fish hosts (Iacuniello *et al*., in prep.) spanning a range of depths (all shallower than 1000 m) and feeding strategies (15, 39). The abyssal and hadal microbial communities were also distinct from those within the broader fish gut dataset, while bathydemersal *Careproctus melanurus* gut communities were interspersed with samples from other shallower fishes (**Figure S2;** species type, R^2^ = 0.46, F=4.25, DF = 16, p < .001).

**Figure 2.**
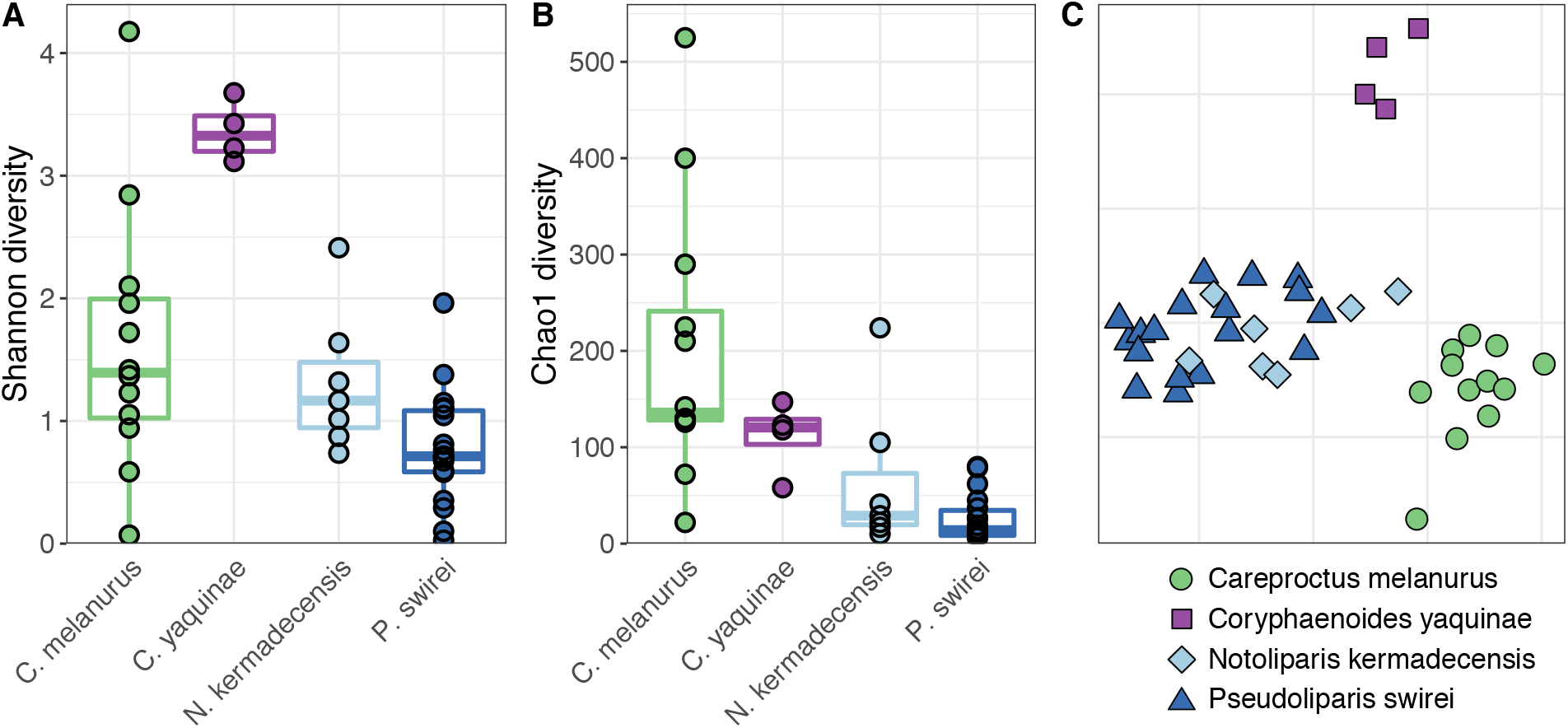
Alpha (A, Shannon; B, Chao1) and beta (C, NMDS ordination based on Bray-Curtis dissimilarity; stress = 0.12) diversity comparisons of the four fishes in this study show that their gut microbiomes are unique from one another. Snailfishes from the continental slope (*Careproctus melanurus*) and hadal trenches (*Notoliparis kermadecensis* and *Pseudoliparis swirei*) are compared to an abyssal rattail (*Coryphaenoides yaquinae*). Colors are the same in all panels, with each species in panel C also reflected by a different shape.

### The hadal snailfish microbiome

The microbiome of *Notoliparis kermadecensis* was dominated by only a few ASVs such that the top ten most abundant sequences made up more than 75% of the communities of each fish (**Figure 3**). Two of the most abundant ASVs were related to the Mycoplasmataceae and composed ∼60% (range ∼ 5–99%) of the gut community. One of these *Mycoplasmataceae* ASVs was identical to the 16S rRNA gene reported from a hadal fish from the Mariana Trench (34) and was distantly related (<95% similar) to sequences found within other fishes (40, 41, 42; **Figure S3**). The second ASV was also similar to those found in other cold-water fish, including notothenioids from Antarctica (43) and grayling from Siberia (44). Together, two ASVs related to *Moritella* made up ∼19% of each community (range ∼0–60%) and were more than 97% similar to 16S rRNA genes from both piezophilic (45) and non-piezophilic taxa. An ASV related to the Desulfovibrionaceae was present in all seven *N. kermadecensis* specimens (mean ∼3%, range ∼0.005–14%). This ASV was highly similar to sequences from notothenioid fish from Antarctic waters (43) and freshwater grayling from Lake Baikal, but less than 95% similar to other sequences (**Figure S4**). We also identified an ASV related to the Rhodobacteraceae (range ∼0–43%) which was present in one sample in high abundance and was identical to sequences from the deep ocean, including the Japan Trench at 7000 m (46). Other abundant ASVs included those classified as members of the *Pseudarthrobacter* (Micrococcaceae; mean ∼5%, range ∼0–17%, identified in every specimen), Corynebacteriaceae (mean ∼0.5%, range ∼0–3%), and *Photobacterium* (average ∼0.5%, range ∼0–1.5%).

**Figure 3.**
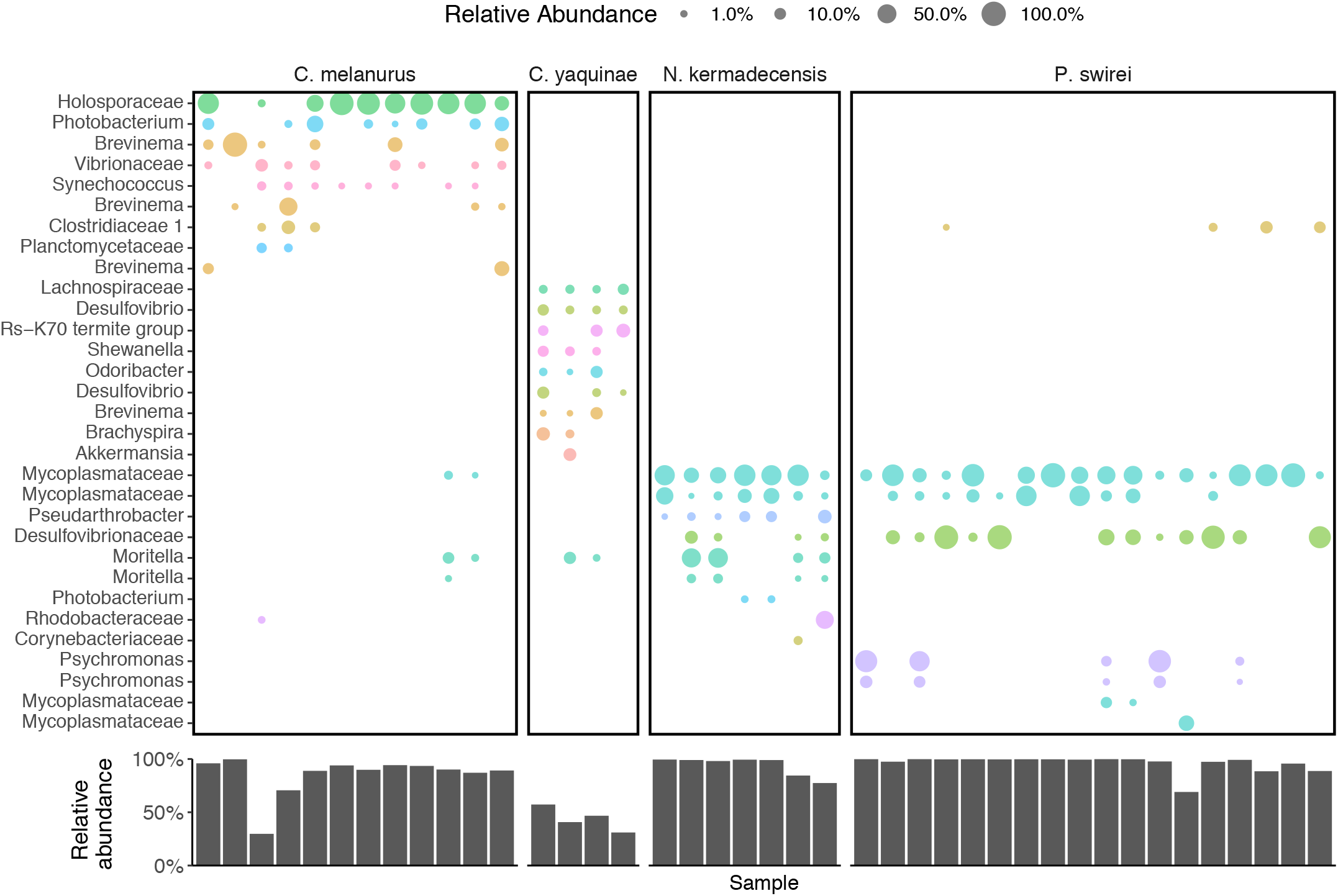
Top; The most abundant ASVs present within each fish species, colored and labeled by their lowest identifiable taxonomic rank. ASVs are shown only if they reach relative abundances greater than 0.5 % in a given sample. Bottom; The total, summed relative abundance of the taxa shown above within each sample.

Like *N. kermadecensis*, the microbiome of *Pseudoliparis swirei* was primarily composed of only a few taxa (**Figure 3**). Members of the Mycoplasmataceae were some of the most abundant (combined mean abundance of four ASVs 53%, range ∼2–99%). Two of these Mycoplasmataceae ASVs were the same as those present in *N. kermadecensis*. A third ASV was present in only one fish but made up ∼27% of that community. The fourth Mycoplasmataceae ASV, closely related to sequences from stone flounder and turbot, was detected in five *P. swirei* specimens with a mean abundance of ∼0.5% (range ∼0-9%). The Desulfovibrionaceae ASV found in *N. kermadecensis* was also present at high abundances in *P. swirei* (mean ∼27%, range ∼0–98%). Other taxa included two ASVs related to the genus *Psychromonas* (combined mean abundance ∼15%, range ∼0–93%). These sequences were similar to known piezophilic microbes obtained from deep-sea amphipod material (**Figure S5;** 7, 47, 48).

We compared the microbial communities in the two hadal fishes against one another. *Psychromonas* was more abundant in the Mariana snailfish, while *Moritella, Pseudarthrobacter*, and *Photobacterium* were enriched in the Kermadec snailfish (**Figure 4C)**. Sequences related to the Mycoplasmataceae and Desulfovibrionaceae were not differentially enriched within either fish.

**Figure 4.**
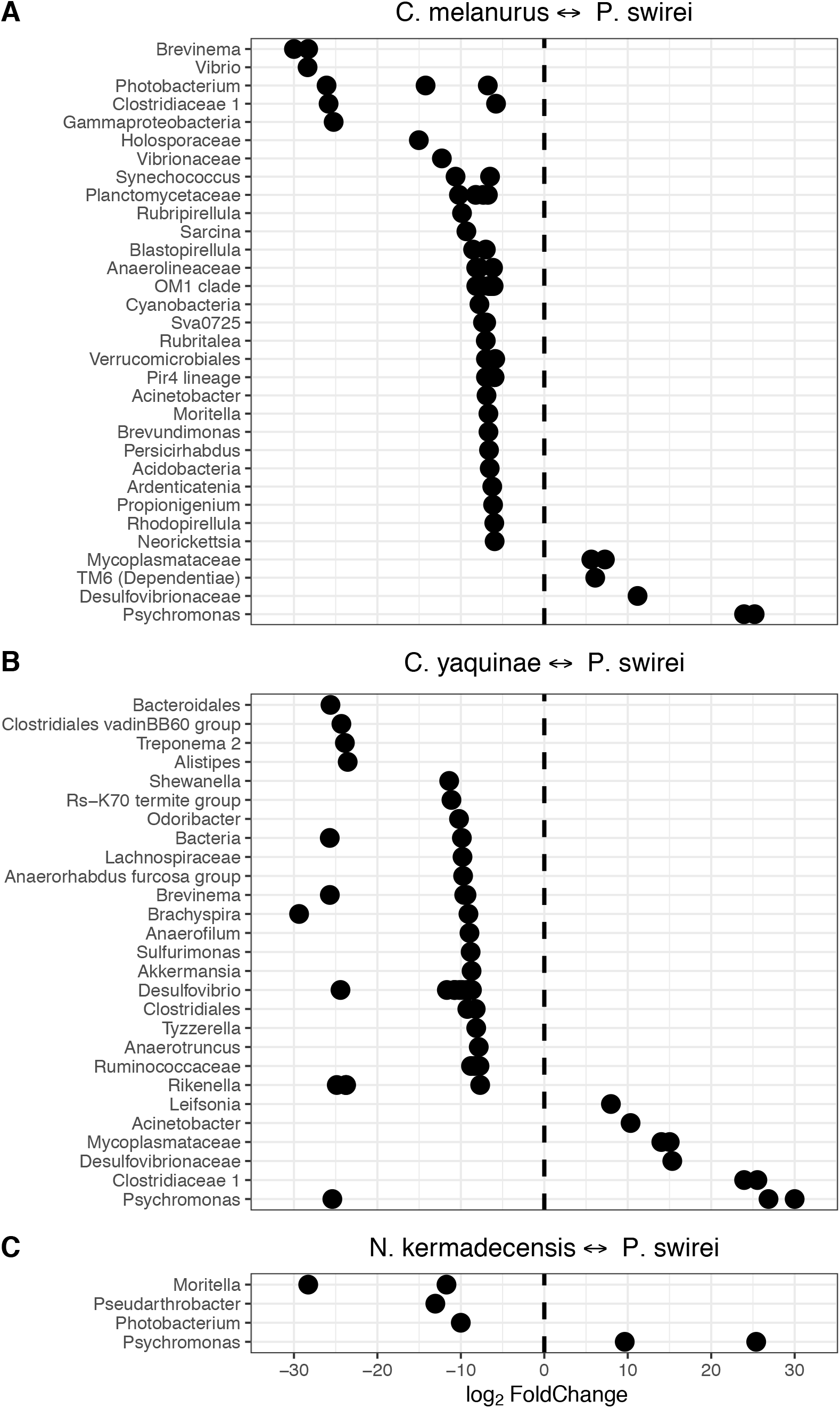
ASVs identified as differentially abundant when comparing fish species against one other. Communities from the Mariana snailfish, *Pseudoliparis swirei*, compared to A) the snailfish *Careproctus melanurus* from the continental slope, B) the abyssal rattail *Coryphaenoides yaquinae*, and C) a hadal snailfish from the Kermadec Trench, *Notoliparis kermadecensis*. ASVs are labeled based on their lowest identifiable taxonomic rank.

### Bathyal, abyssal, and hadal fish gut microbiome comparisons

We also analyzed the microbiota of the shallower-living snailfish *Careproctus melanurus* collected from ∼300–800 m depth. Although this fish had higher alpha diversity than the hadal snailfishes (**Figure 2**), the gut-associated microbial community was still dominated by only a handful of sequences (**Figure 3**). The most abundant ASV was related to the Holosporaceae, (mean ∼50%, range ∼0–92%) and showed >97% sequence similarity to taxa identified within other host-associated systems (49). Other ASVs included three relatives of the genus *Brevinema* (combined mean abundance ∼20%, range ∼0–100%) which were similar to those found in graylings from Lake Baikal, mudsuckers, and unicornfish. We note the presence of two ASVs related to the genera *Moritella* (combined mean ∼1%, range 0–11%) and *Photobacterium* (combined mean ∼7%, range ∼0–33%). These ASVs were similar (>99%) to both known piezophilic and piezosensitive species. Other abundant ASVs included taxa in the *Vibrionaceae* (mean ∼3%, range ∼0–13%), Clostridiaceae (mean ∼2%, range ∼0–17%), and *Synechococcus* (mean ∼1%, range ∼0.005– 4%). The Clostridiaceae ASV was present in both *N. kermadecensis* and *P. swirei* at low abundances.

When comparing *C. melanurus* against *P. swirei*, many ASVs were more abundant in the continental slope-dwelling fish, reflecting the overall lower alpha diversity of the hadal snailfish (**Figure 4**). Amongst the Gammaproteobacteria, sequences related to the genera *Vibrio, Moritella*, and *Photobacterium* were more abundant in *C. melanurus*, while *Psychromonas* was more abundant in *P. swirei*. Other taxa of note included the enrichment of *Synechococcus* and other Cyanobacteria within the shallower fish, and the enrichment of ASVs related to the Mycoplasmataceae, Desulfovibrionaceae, and the phylum TM6 within *P. swirei*. The sequence belonging to the phylum TM6 was similar to those collected from deep-ocean sediments (50, 51). Comparisons between *C. melanurus* and *N. kermadecensis* revealed similar trends in differentially abundant taxa as with *P. swirei*.

In contrast to the snailfishes, microbial community composition within the gut of the rattail *Coryphaenoides yaquinae*, collected from 4000–6000 m depth, was much more even. The ten most abundant ASVs represented 31–57% of the community (**Figure 3**). Four of the top ten most abundant ASVs, related to the *Desulfovibrio*, Deltaproteobacteria group Rs-K70, *Brachyspira*, and family Lachnospiraceae (combined mean ∼22%, range 6-37%), were most closely related to sequences from various low-oxygen environments. A further four ASVs had highest identity to sequences from fish samples and were classified as belonging to the genera *Akkermansia, Brevinema, Desulfovibrio*, and *Odoribacter* (combined mean ∼15%, range 3-27%). We also identified sequences similar to *Shewanella* (mean ∼4%, range ∼0.2–8%) and *Moritella* (mean ∼3.5%, range ∼0.3–12%) in all *Coryphaenoides yaquinae* specimens. The ASV related to *Shewanella* was most similar to the piezophiles *S. benthica* KT99 and *S. violacea* (52, 53) and to sequences previously identified from *Coryphaenoides yaquinae* (**Figure S6**; 6). The *Moritella* ASV was the same as that within *N. kermadecensis* and *C. melanurus* and was highly similar (>99%) to both piezophilic and piezosensitive strains. Because the communities of *C. yaquinae* and *P. swirei* were so distinct from one another, comparisons between the two fishes showed that many of the differentially abundant taxa were also the dominant members of the respective communities (**Figure 4**).

Finally, we leveraged a broader dataset of fishes and environmental samples to investigate microbial lineages specific to the hadal fishes. We first screened these samples for specific ASVs that were abundant in the hadal fishes, including those related to the Mycoplasmataceae, Desulfovibrionaceae, *Psychromonas, Moritella*, and *Shewanella*. While these lineages dominated the hadal samples, they were not found at high abundances in any other fish (**Figure S7**). These ASVs represented a miniscule fraction of Mariana Trench sediment and water samples, reflecting on average only 0.007% of the community (identified in four of 16 samples; maximum abundance, 0.036%). We broadened our search to include any ASV related to the Mycoplasmataceae and found that many of the shallower fish gut microbiomes contained this family (**Figure S8)**. While we did not find high abundances of sequences related to known piezophilic lineages in the comparison fishes, we found that almost all of the shallower-living fish gut communities included *Photobacterium* and *Vibrio* (**Figure 5**).

**Figure 5.**
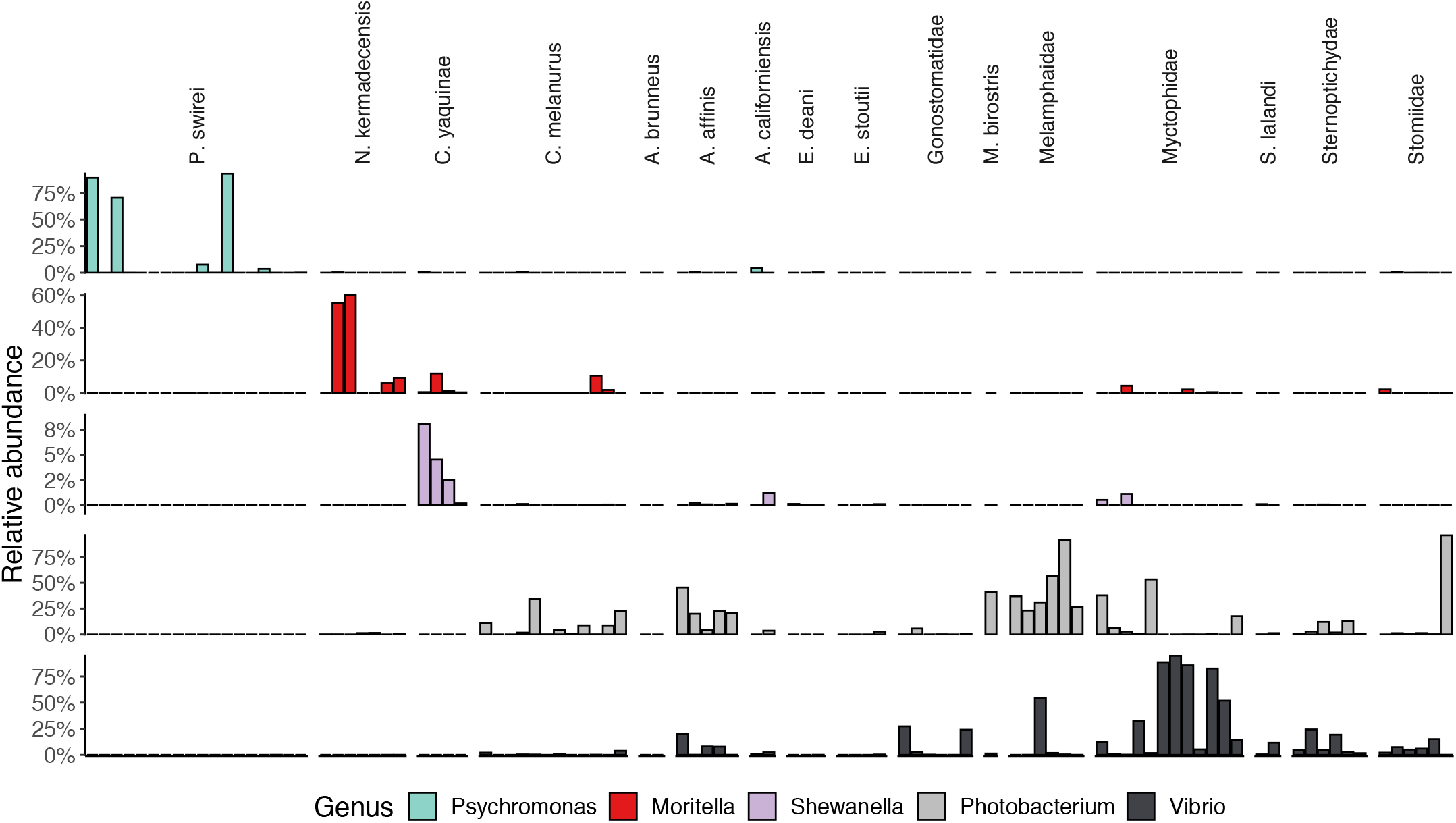
The abundances of gammaproteobacterial genera which are known to contain cultured piezophilic and/or piezosensitive members within the four comparison species and a wider dataset of fishes. (Top row, *Psychromonas*; second row, *Moritella*; third row; *Shewanella*; fourth row, *Photobacterium*; bottom row; *Vibrio*).

## Discussion

We describe the gut-associated microbial communities within the hadal snailfishes *Pseudoliparis swirei* and *Notoliparis kermadecensis*, the abyssal rattail *Coryphaenoides yaquinae*, and the continental slope-dwelling snailfish *Careproctus melanurus*. These fishes include some of the dominant vertebrates at abyssal and hadal depths. The microbial communities within these four species were distinct from one another. Our findings show that while the shallow and deep-water snailfishes belong to the same family, they have large differences in their gut microbiota. The communities were also different between the abyssal and hadal teleosts, indicating fishes that experience similar environmental conditions at great depth do not necessarily have similar microbial gut flora. In contrast, many of the most abundant lineages were shared between both hadal snailfishes. It has been proposed that *Pseudoliparis* and *Notoliparis* should be synonymized as one genus (24, 54), suggesting these fish may have similar host physico-chemical variables that could influence their microbiome (e.g. pH, O2; 55, 56, 57). One explanation for the observed differences in the fish gut microbiomes could be diet. Host trophic strategy influences the diversity of microbial communities within the gut (56, 57, 58, 59) and may be one reason for the shift in fishes at the abyssal-hadal boundary (60). Hadal snailfishes primarily eat amphipods and occasionally polychaetes and decapod shrimp, reflecting a restrictive diet. The diet of *Careproctus melanurus* also consists of crustaceans, including amphipods, shrimp, mysids, and tanaids, but can also include bivalves, polychaetes, and fish (61, 62). In contrast, the diet of *C. yaquinae* is composed primarily of carrion, along with squid, crustaceans, and other fish (18, 33). The relatively narrow dietary choices of hadal fishes may shape their gut microbiomes in relation to shallower fishes. Future work should investigate how host physiology, diet, and environmental factors, such as differences in water mass or organic matter input (32, 63, 64), impact deep-sea fish gut microbiota.

The hadal fish gut microbiomes were composed of only a few ASVs and largely dominated by members of the Mycoplasmataceae, a family common in the digestive tracts of fishes (41, 65). While *Mycoplasma* can be pathogenic (66, 67, 68), it was recently suggested that Mycoplasmataceae in *P. swirei* may supply the host with the cofactor riboflavin (34). Based on our analyses, there are multiple strains of Mycoplasmataceae present within hadal snailfishes. This family was present in nearly all fishes in the broader dataset but was not as abundant in our two comparison species. Instead, *C. melanurus* appears to have high abundances of the Holosporaceae, a group which are known to infect shrimp and ciliates (49, 69). We show that known host-associated, potentially-pathogenic lineages such as Mycoplasmataceae are common in fishes from the surface ocean to hadal depths, although with apparent differences at the ASV level. The Mycoplasmataceae represent interesting targets for identifying adaptations to high pressure because of their exceptionally reduced genome sizes and limited metabolic functionality (70).

Trenches are typically isolated by large expanses where seafloor depths are shallower than 6,000 m. If we assume that hadal species are obligately adapted to *in situ* pressures, trenches would have high rates of biogeographic isolation. Indeed, many megafaunal species found in trenches appear to be endemic (32, 71, 72), including hadal snailfishes which appear genetically isolated from one another (24). Despite the geographic and genetic separation of the hosts, several identical ASVs were found within both hadal snailfishes, including those related to the Mycoplasmataceae and Desulfovibrionaceae. Neither the Mycoplasmatacaeae nor Desulfovibrionaceae sequences were present in high abundances in any of the other fishes analyzed in this study. One explanation is that the extant Mycoplasmataceae have not undergone appreciable genomic evolution to diverge from the ancestral symbiont present within snailfishes prior to their radiation into separate trenches approximately 20–40 mya (54). An alternative explanation could be the dispersal of very closely related microbial taxa between two trenches 6,000 km apart and subsequent host selection for these lineages. Certain microbial symbionts are highly specific within deep-sea anglerfishes and may be dispersed horizontally through the water column (73, 74). The possibility of dispersal of water and sediment microorganisms between trenches has been previously highlighted (12, 13, 75). Whole genome sequencing, e.g. metagenome-assembled genomes, will be required to determine if these strains are similar beyond their 16S rRNA gene and to understand their dispersal and evolution in hadal habitats.

We show that abyssal and hadal fishes have high abundances of sequences related to known piezophilic taxa, including *Psychromonas, Moritella*, and *Shewanella*. This finding is consistent with the observation that guts of deep-sea animals can show high levels of piezophily (5, 76, 77, 78) and that piezophiles are most successfully cultivated from deep-sea hosts (6, 7, 8, 52, 79, 80). This is in contrast to hadal water and sediment communities where sequences associated with previously cultured piezophiles represent relative abundances of less than 1% (11, 12, 13, 81). We therefore add to a growing body of evidence that known, isolated piezophilic genera are associated with deep-sea animals. However, different piezophilic lineages were abundant in each species of fish: *Psychromonas* in *P. swirei, Moritella* in *N. kermadecensis*, and *Shewanella* in *C. yaquinae*. None of the Mariana snailfish reported in a different study had high abundances of *Psychromonas* (n=2; 34). One hypothesis is that these piezophilic taxa may represent more transient members of the fish gut, for example acquired through the consumption of amphipods (e.g. *Psychromonas*; 48, 82, 83, 84). If there was a strong signal from transient taxa, we might expect to see *Pseudoalteromonas* or *Psychrobacter*, which can reach abundances of >20% of the gut-associated microbiota of amphipods in the Mariana Trench (83, 84), within the guts of amphipod-feeding hadal fish. However, we did not find these genera in appreciable abundances in any of the abyssal or hadal fish (maximum of 0.13% for *Pseudoalteromonas* and 0.36% for *Psychrobacter*). It is therefore likely that the piezophilic microbes are present at least in part because of host-microbe specificity. Indeed, piezophilic *Colwellia* with > 99% average genomic nucleotide identity have been isolated from deep-sea amphipods collected over 30 years apart (85), highlighting strong selection temporally in some hadal organisms. Representative piezophiles can contain genes encoding chitinase, including isolates belonging to *Psychromonas, Moritella, Shewanella*, and *Colwellia* (53, 80, 85, 86), and sediments amended with chitin also showed a response of known piezophilic taxa (87). Moreover, a recent metagenomic analysis of salmonid fishes revealed gut-associated *Mycoplasma* harbor genes putatively involved in the degradation of long-chain polymers such as chitin (88). Amendments of fish guts with chitin, coupled to metagenomic and transcriptomic sequencing, may reveal catabolic functions that benefit the host via the processing of recalcitrant dietary compounds.

One unifying characteristic of these piezophilic Gammaproteobacteria is the synthesis of long-chain omega-3 polyunsaturated fatty acids (LC-PUFAs), with *Psychromonas, Moritella*, and *Colwellia* species producing docosahexaenoic acid (DHA, 22:6*n-*3) and *Shewanella* species producing eicosapentaenoic acid (EPA, 20:5*n-*3; 89). LC-PUFAs are essential fatty acids required for proper development and growth of all metazoans yet most vertebrates are unable to synthesize them *de novo*, thus, they need to be obtained from the diet. In shallow marine habitats, phytoplankton are the primary producers of LC-PUFAs however the quality and quantity of these essential fatty acids that reach abyssal and hadal zones is minimal. It is thus compelling to hypothesize that the enrichment of LC-PUFA producing taxa in hadal metazoan microbiomes may represent the primary source for delivery of these essential fatty acid nutrients to their hosts.

While members of piezophilic genera were not present in the broader fish dataset analyzed here, we instead found high abundances of the gammaproteobacterial genera *Photobacterium* and *Vibrio* (**Figure 5**). *Photobacterium* are common within microbiomes of marine fishes (90, 91, 92) and can be moderate piezophiles, with some strains showing growth up to 70 MPa (93, 94). To our knowledge, no member of the genus *Photobacterium* has been isolated at *in situ* pressures from hadal depths. Although the *Photobacterium* ASVs in *C. melanurus* and *N. kermadecensis* are distinct, the high similarity of the 16S rRNA genes of piezophilic and non-piezophilic ecotypes (95) precludes an analysis here of their putative pressure sensitivity. We present the hypothesis that there may be a change in the dominant heterotrophic Gammaproteobacteria within the microbiomes of animals as a result of the selective pressure of increasing water depth. At shallower depths (e.g. 0 – 2000 m), taxa such as *Photobacterium* and *Vibrio* may be abundant, but with increasing depth the gut community may shift towards hyperpiezophiles, including members of the genera *Psychromonas, Moritella*, and *Shewanella*. An analysis of fish gut microbial communities along a more comprehensive depth gradient, for example targeting depths between 1000 – 4000 m, will be needed to assess this hypothesis.

The observation that representatives of known, isolated piezophilic taxa are abundant within deep-ocean animals reveals two important insights into the lack of high pressure-adapted isolate diversity in the literature. First, nearly all attempts to isolate microbes from abyssal and hadal samples have used nutrient-rich media which ultimately select for copiotrophic lineages. The gut of a host would similarly select for taxa capable of taking advantage of a high-nutrient environment, unlike the carbon-limited niches in deep-ocean water or sediments. Second, pressure vessels are generally static incubation chambers, requiring organisms to cope with variable waste, oxygen, and nutrient concentrations. Similar conditions might be expected in the guts of an abyssal or hadal fish undergoing varying events of feast and starvation. Piezophilic taxa are capable of responding to variable environmental conditions: a non-exhaustive list includes enrichment of these groups on detritus (87), particles (96, 97), oil and dispersant (98, 99, 100, 101, 102), methane (103), under low oxygen conditions (104), in eukaryotic mesocosms (105), in pressure-retaining samplers (106), and in hadal sediments after long-term, static, and unamended conditions (13). Therefore, genera such as *Psychromonas, Shewanella, Moritella*, and *Colwellia* are likely isolated because of their ability to adapt to the variable nutrient and oxygen conditions found both within the guts of deep-sea megafauna and the pressure vessels used for cultivation in the laboratory. The implications of this observation are that static mesocosms performed in the lab using current methods will almost always select for a distinct group of microorganisms that are not representative of environmental deep-sea communities at large, but which nonetheless fill a specific niche in the deep ocean on particles and in the guts of megafauna.

In addition to clarifying the role of recognized, lab-characterized piezophilic lineages, the examination of microbial diversity associated with extreme deep-sea animals reveals new taxa that likely possess pressure-adapted lifestyles. These taxa represent broad phylogenetic groups that significantly extend the hyperpiezophile ranks beyond the Gammaproteobacteria, including Mycoplasmataceae, Desulfobacterota, and Actinobacteria (*Pseudarthrobacter*). For example, the presence of Desulfovibrionaceae ASVs within both abyssal and hadal fishes may indicate the presence of sulfate reduction occurring within the guts of deep-ocean fishes at high hydrostatic pressure. Future studies that integrate metagenomic profiling combined with novel cultivation approaches that mimic the *in vivo* fish gut microbial ecosystem will be required to more fully define the breadth of metabolic activities that support the success of the microbes and the fish they inhabit within the deep sea.

## Methods

### Sample collection

Abyssal and hadal fishes were collected from the Kermadec and Mariana trenches aboard the R/V *Thomas G. Thompson* and R/V *Falkor* during April–May 2014 and November–December 2014, respectively. Fishes were caught using free-vehicle lander systems equipped with acoustic releases (26). The traps were baited with mackerel and squid wrapped in nylon mesh to limit bait ingestion by sampled taxa. Once fish specimens were on board they were immediately placed on ice and processed. Gut material was carefully extracted from the hindgut, flash-frozen in liquid nitrogen either dry or in RNA Later, and stored at -80°C. One *Notoliparis* specimen was reported as belonging to a different species, *Notoliparis stewarti*, based on morphological characteristics (NK100329; 107). Because of the similarity of these two potentially different species, their presentation as the same species in previous publications, and the apparent similarity of their microbiomes, we report this one specimen as *N. kermadecensis* throughout this manuscript but acknowledge future work is needed to fully characterize the taxonomy hadal fishes. Specimens of *Careproctus melanurus* were collected by trawl aboard the F/V *Noah’s Ark* and F/V *Last Straw* during the summer 2014 NOAA NWFSC Groundfish Bottom Trawl Survey. One specimen was also collected from a 2015 UC Ship Funds-supported student cruise aboard the R/V *Sproul*. Following the storage of whole fishes at -20°C, specimens were defrosted and the hindgut dissected. Gut contents from *C. melanurus* were submerged in Chaos lysis buffer (5M guanidine thiocyanate, 2% sarkosyl, 50 mM EDTA, 40 ug/ml proteinase K, and 15% beta-mercaptoethanol) and stored at -80°C prior to analysis. We acknowledge that these slightly different methods of sample processing and preservation, given the constraints of shipboard sample processing, may influence microbial community composition downstream.

### DNA extraction and 16S rRNA gene amplicon sequencing

DNA was extracted from gut samples using an organic extraction method. Intestinal contents were defrosted and resuspended in Chaos buffer. After a 30 min incubation at 55°C, samples were homogenized by bead-beating with silica beads. Lysate was then treated with one volume of phenol:chloroform:isoamyl alcohol (25:24:1). DNA in the resulting aqueous layer was cleaned with the Zymo Research Quick-gDNA MiniPrep kit (Irvine, CA). Negative control extractions were performed with each set of fish samples.

After extraction, the V4 region (∼290 bp) of the 16S rRNA gene was amplified using a two-step PCR protocol to create dual-barcoded amplicons. The first reaction used primers 515F-Y and 806rb with overhangs for attachment of Illumina-compatible indexes in the second reaction (108). The initial reaction was performed in triplicate using Q5 polymerase (NEB, Ipswitch, MA) as follows: initial denaturation of 30 s at 98°C; 25 cycles of 10 s at 98°C, 20 s at 50°C, 30 s at 72°C; final extension of 2 min at 72°C. Triplicate reactions were combined and 5 μL of each sample pool was used as template in a second reaction to attach unique indexing primer pairs. The second reaction was performed as above except using only 8 cycles and an annealing temperature of 56°C. Barcoded amplicons were cleaned using AMPure XP Beads (Beckman Coulter, Brea, CA), pooled in equimolar concentrations, and sequenced on Illumina’s MiSeq platform (2 × 300 bp) at the UC San Diego Institute for Genomic Medicine and the UC Davis Genome Center.

### Sequence processing and analysis

Paired raw reads were trimmed with Trimmomatic v0.35 (109) and filtered to sequences ≥100 bp. Trimmed reads were imported into the QIIME 2 platform v2018.6 (110) where the Dada2 workflow plugin v2018.6 (111) was used to trim primer regions, denoise, and merge sequences to generate ASVs. Chimeras were removed using the consensus method. Non-ribosomal sequences were excluded and taxonomy was assigned to ASVs using the scikit-learn naive Bayes machine-learning classifier (112) in QIIME 2 trained on the SILVA v128 SSU database (113). Further filtering and all downstream analyses were performed in R (114). Singletons, sequences classified as eukaryotic, or those unassigned at the domain level were removed. To ensure a conservative analysis, potential contaminants were identified by co-occurrence network of all ASVs using the R package *ccrepe* v1.24.0 (115). Twelve ASVs which belonged to the genera *Acinetobacter* and *Pseudomonas* and the families Comamonadaceae, Caulobacteraceae, and Methylophilaceae were identified as both co-occurring and representative of common contaminants (116). These ASVs were filtered from all samples (**Figure S9**). Alpha and beta diversity of communities were estimated using the R package *phyloseq* v1.32.0 (117). Differentially abundant taxa between the different fishes were identified using *DESeq2* v1.28.1 (118) with ASVs of less than 10 reads excluded. We statistically tested the importance of host species on structuring Bray-Curtis dissimilarity by permutational analysis of variance (PERMANOVA) using the adonis and pairwiseAdonis functions (119, 120). For phylogenetic analyses, representative 16S rRNA gene sequences were aligned using the SINA Aligner (121) and trees built using FastTree using default settings (122). Trees were visualized using the Interactive Tree of Life (iTOL; 123).

For further context, the gut microbiomes of the four fish of interest were compared to microbial datasets from a diverse collection of fishes from depths shallower than 1000 m and water and sediments from the Mariana Trench. The comparative fish dataset included catshark (*Apristurus brunneus*), hatchetfish (*Argyropelecus affinis* and other Sternoptichydae), smelt (*Atherinopsis californiensis*), hagfish (*Eptatretus deani, Eptatretus stoutii*), bristlemouths (Gonostomatidae), ridgehead (Melamphaidae), manta ray feces (*Mobula birostris*), lanternfish (Myctophidae), California yellowtail (*Seriola lalandi*), and dragonfishes (Stomiidae). These fishes were typically frozen at -20°C prior to hindgut dissection and then processed in the same manner as described above. The complete microbial communities of these fish will be described elsewhere (Iacuniello *et al*., in prep). The water (RG02, RG07, RG08, RG16, RG18; 3.0, 0.2, and 0.1 μm size-fractionated samples) and sediment samples (FVCR02, FVCR03, FVCR04; 0-1 cm depth fraction) were collected from depths exceeding 5,000 m in the Mariana Trench. A full description of their collection and extraction has been previously described (12, 13). PCR amplification and all further downstream analyses were performed as described above.

## Data Availability

Raw sequencing data for the fish species in this study have been submitted to the NCBI Short Read Archive under BioProject PRJNA720542.

## Acknowledgements

We thank the crews of the R/V *Falkor*, R/V *Thompson*, R/V *Sproul*, F/V *Noah’s Ark*, and F/V *Last Straw* for help at sea. We are grateful to all members of the HADal Ecosystem Studies (HADES) team for scientific advice. We would like to express our appreciation for the financial support provided by the National Science Foundation (1130712 to JCD, 1536776 to DHB, MCB-114552 and OCE-1837116 to EEA), the Schmidt Ocean Institute (cruise FK141109), and the Prince Albert II Foundation (Project 1265 to DHB).

