## Supplemental Figures for "Microbiomes of hadal fishes contain similar taxa, obligate symbionts, and known piezophiles across trench habitats"

**Supplementary Figure 1**. Chao1 (A) and Shannon (B) alpha diversity of the different fishes sequenced in this study, including the wider dataset. The diversity of the wider dataset will be described elsewhere (Iacuaniello *et al*., in prep).

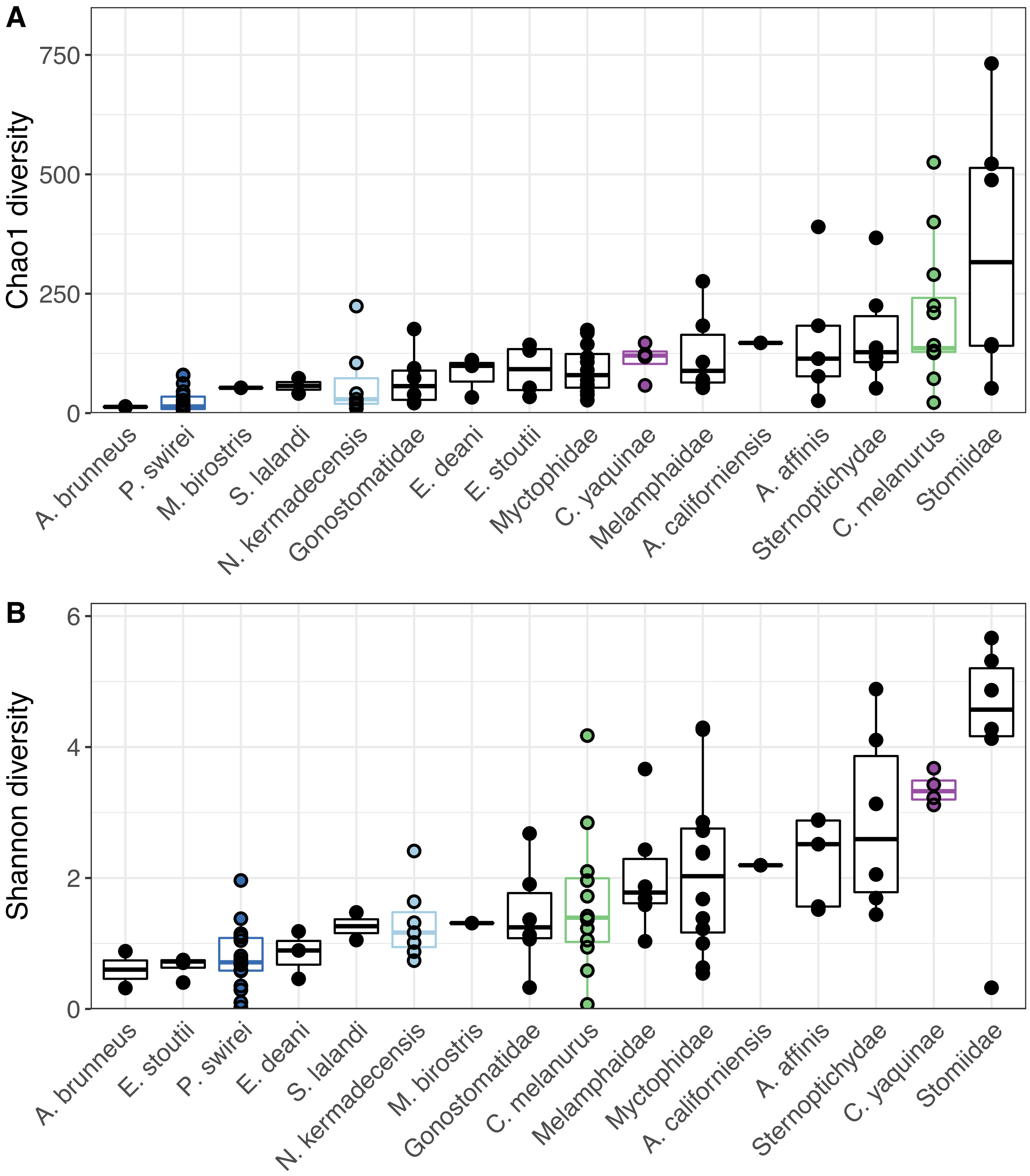

**Supplementary Figure 2**. NMDS ordination based on Bray-Curtis beta diversity of the four study fishes and a wider dataset as described in the text (stress = 20.7). The diversity of the wider dataset will be described elsewhere (Iacuaniello *et al*., in prep) and those samples are shown as empty circles.

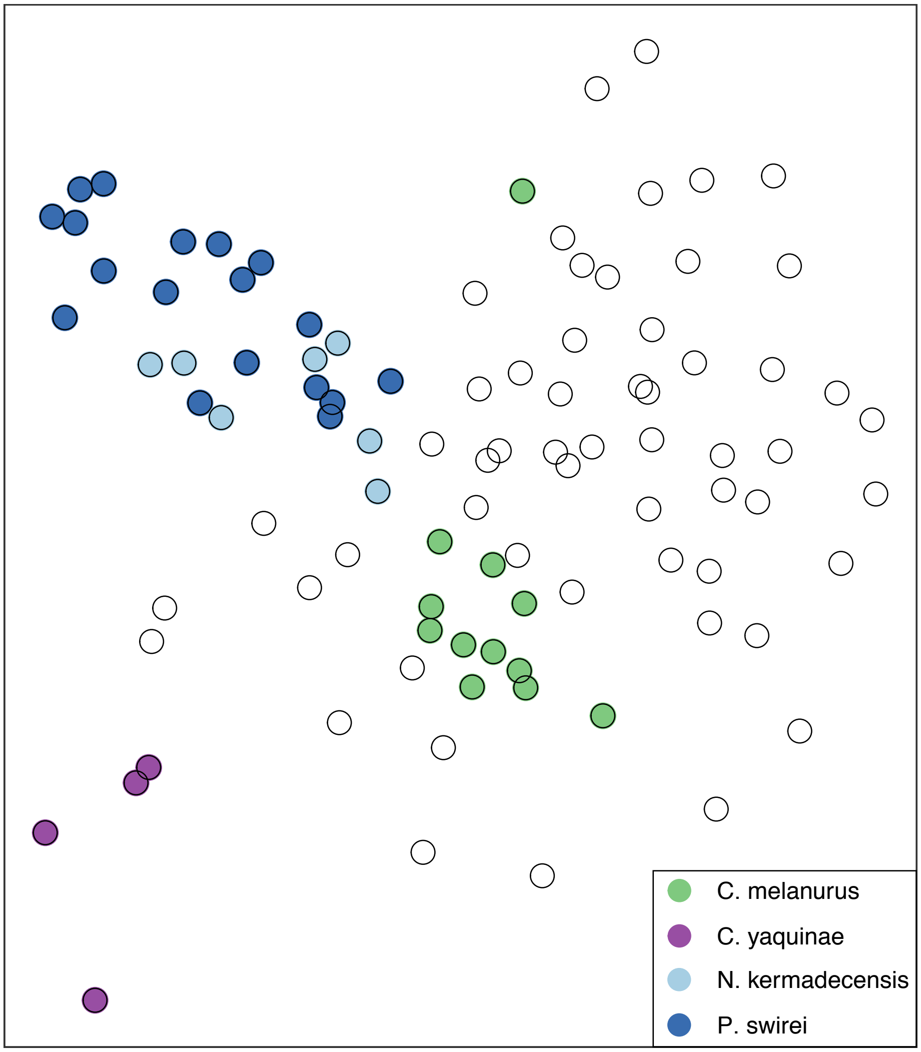

**Supplementary Figure 3**. Phylogenetic tree of Mycoplasmataceae ASVs within the Mariana and Kermadec snailfish. The sequences from this study are shown in blue.

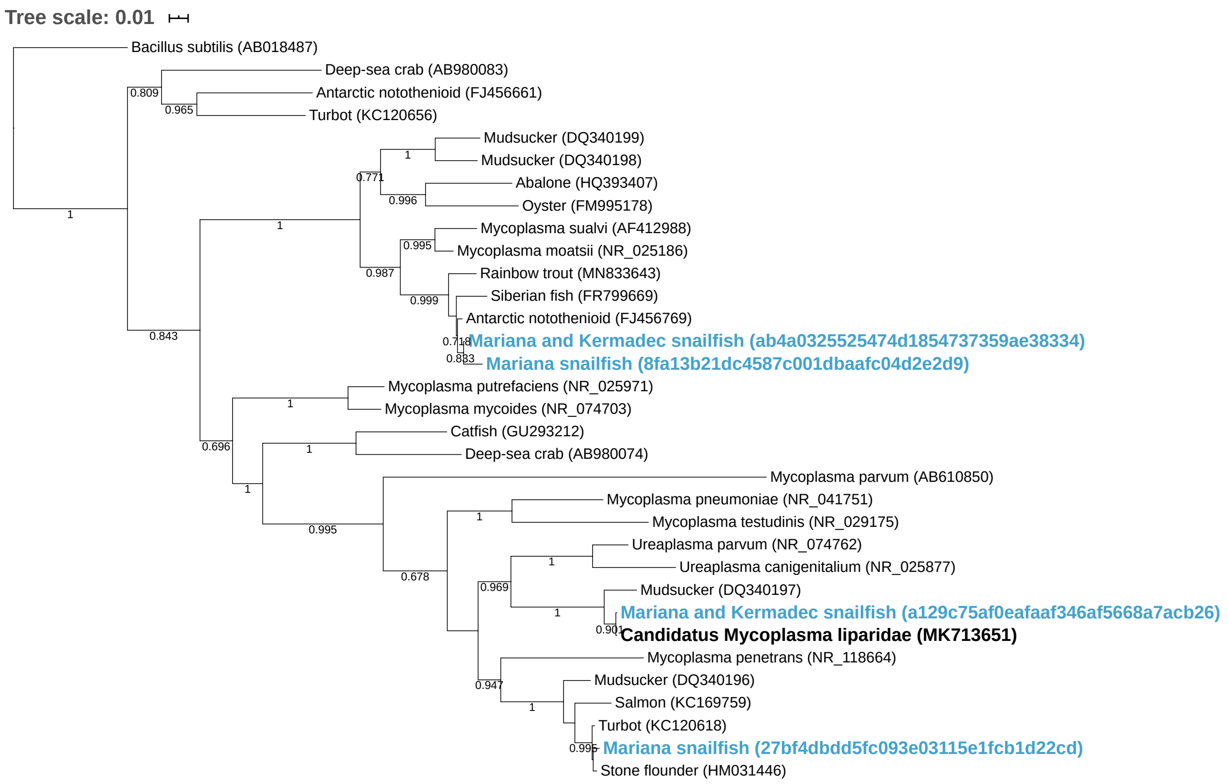

**Supplementary Figure 4.** Phylogenetic tree of an ASV related to the Desulfovibrionaceae found in both the Mariana and Kermadec snailfish. The sequence from this study is shown in blue.

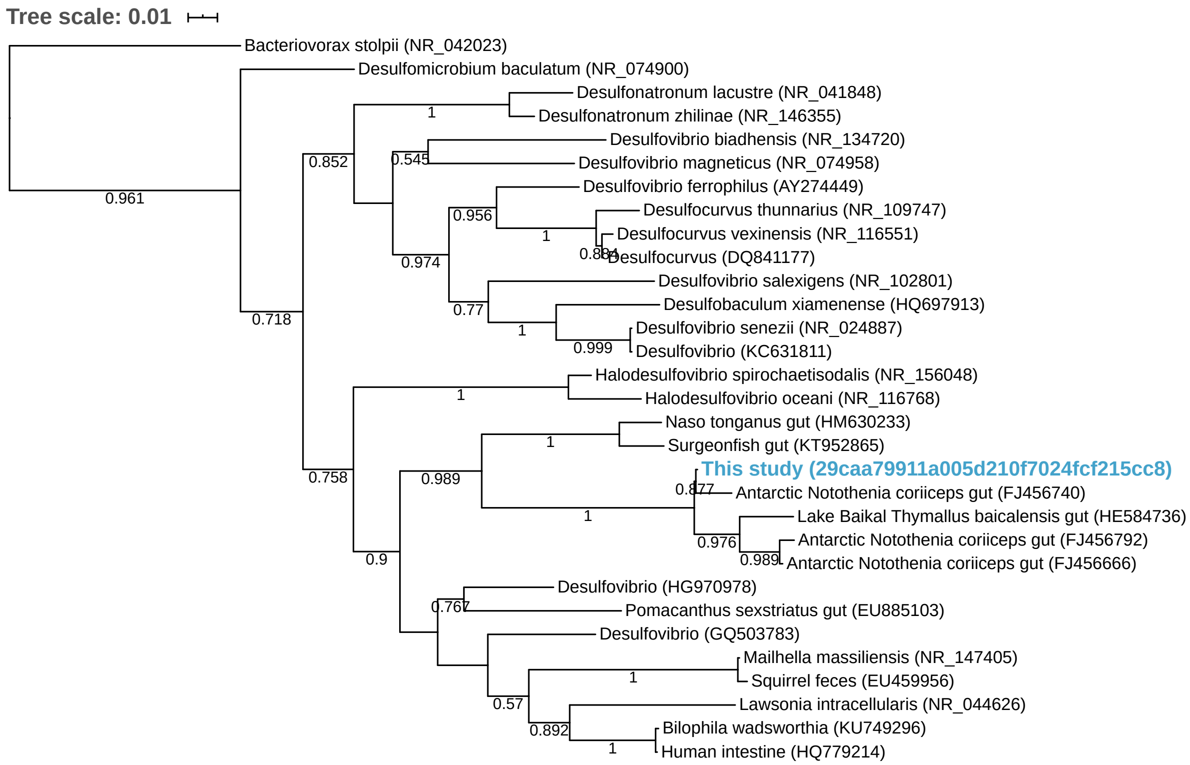

**Supplementary Figure 5**. Phylogenetic tree of two *Psychromonas* ASVs abundant in *P. swirei*. Sequences in this study are shown in blue, while those in black bold are either known piezophiles or obtained from abyssal or hadal environments.

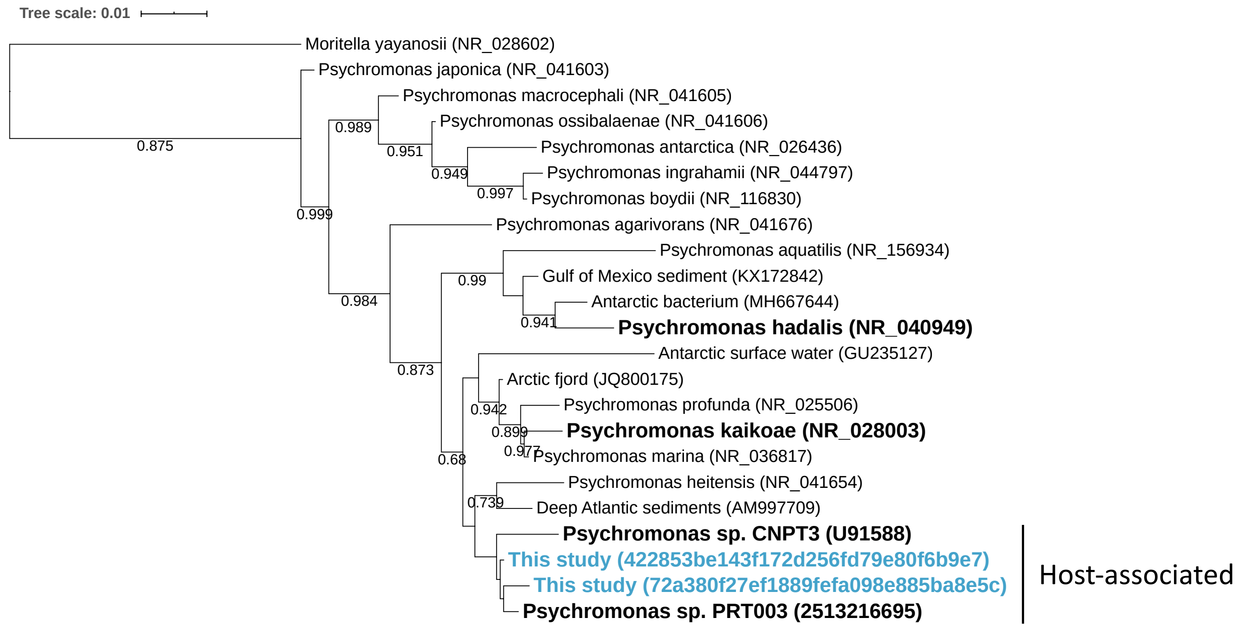

**Supplementary Figure 6**. Phylogenetic tree of an ASV related to *Shewanella* present within *Coryphaenoides yaquinae*. Sequences in this study are shown in blue, while those in black bold are known piezophiles.

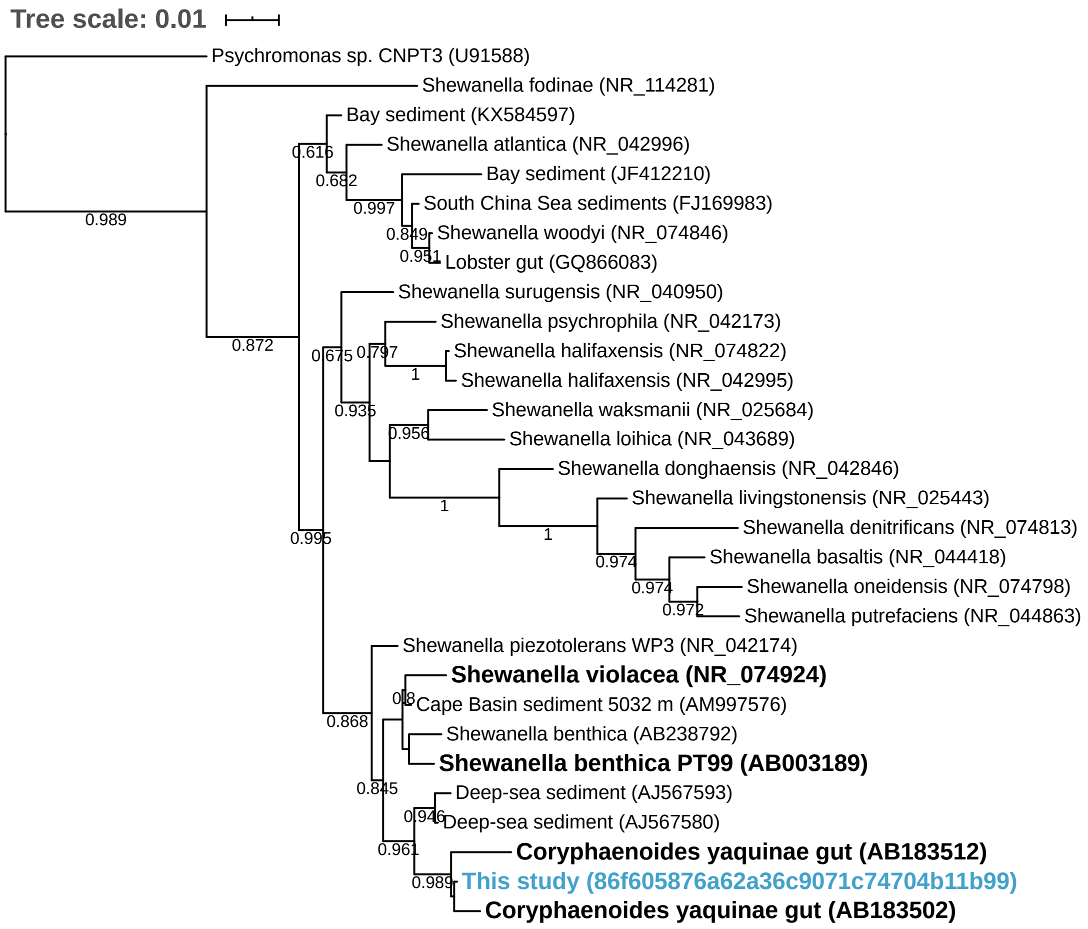

**Supplementary Figure 7.** The presence of seven hadal-abundant ASVs within the four comparison species and a wider dataset of fishes. ASVs are labeled based on their lowest identifiable taxonomic rank.

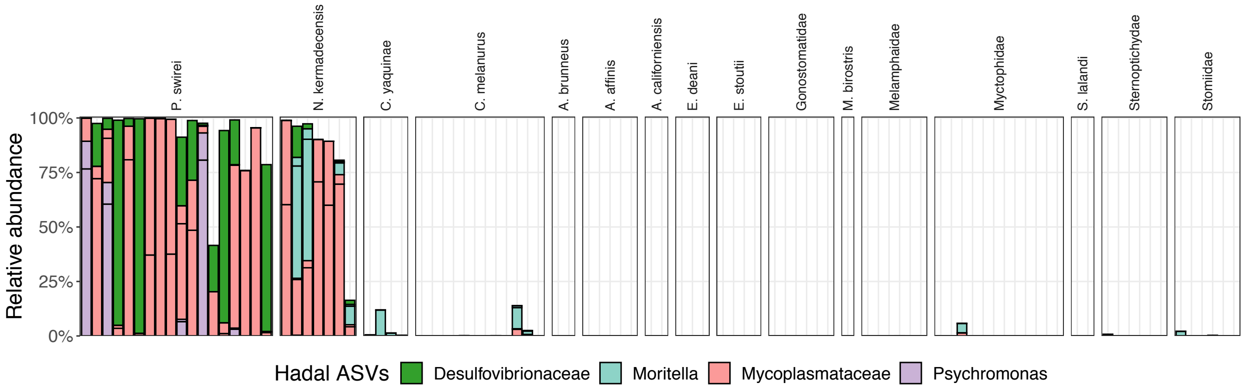

**Supplementary Figure 8**. Relative abundances of ASVs related to the family Mycoplasmataceae within a broad dataset of fish species.

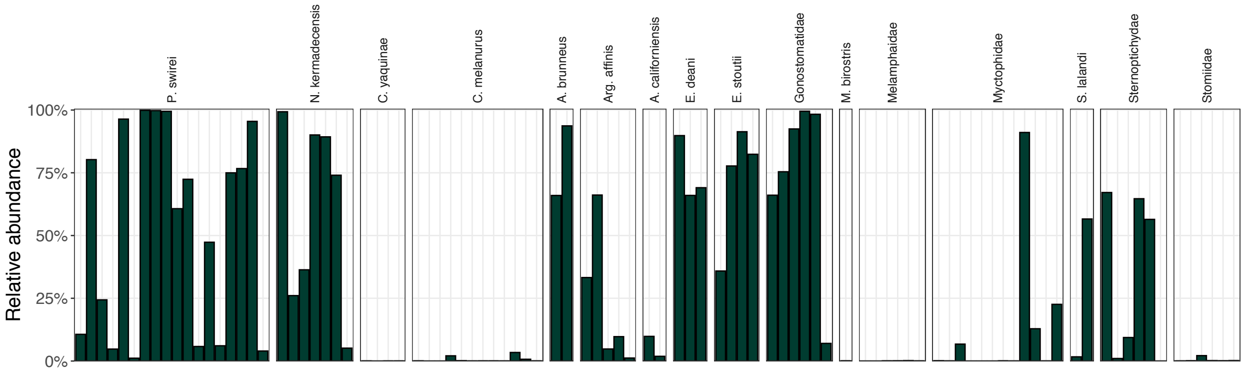

**Supplementary Figure 9**. Co-occurrence map highlighting clusters of taxa common in low-biomass samples, potentially representing contamination sequences. Analysis was performed using CCREPE, with 12 taxa identified and filtered from libraries before downstream analysis.

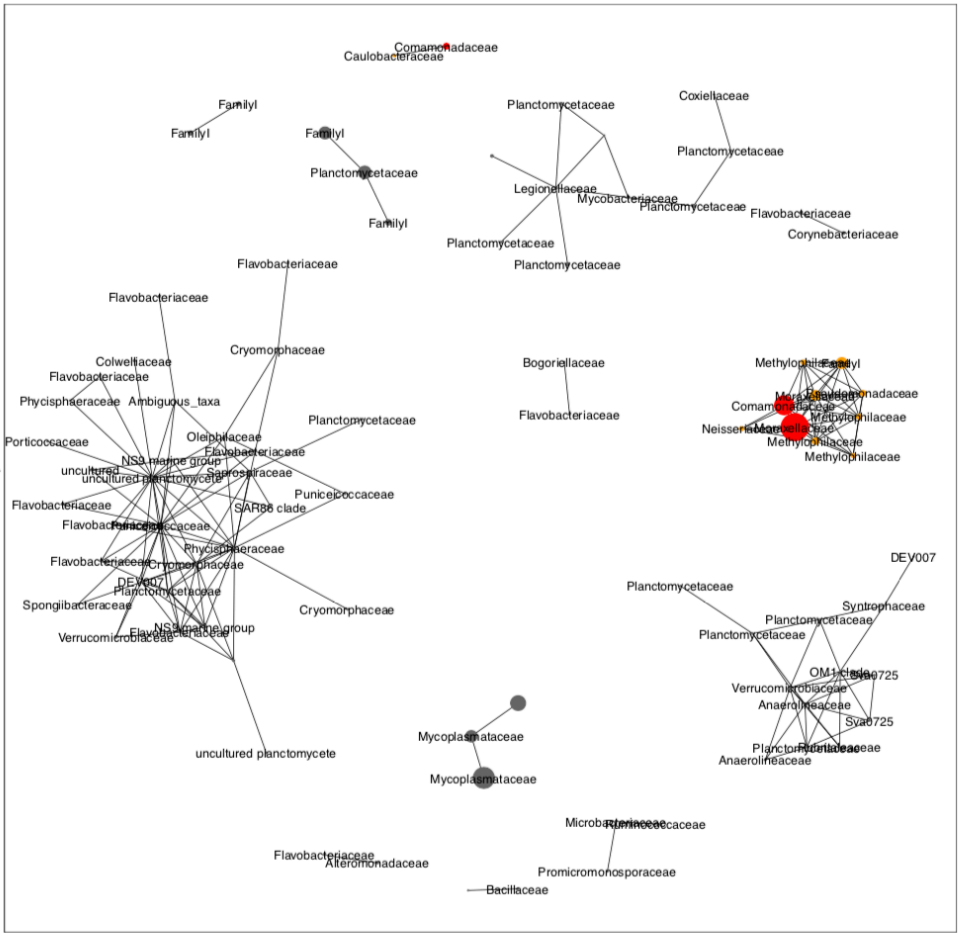
